# Nutritional status and fecundity are synchronised by muscular exopheresis

**DOI:** 10.1101/2020.06.17.157230

**Authors:** Michał Turek, Małgorzata Piechota, Katarzyna Banasiak, Nilesh Shanmugam, Matylda Macias, Małgorzata Alicja Śliwińska, Marta Niklewicz, Konrad Kowalski, Natalia Nowak, Agnieszka Chacińska, Wojciech Pokrzywa

**Author notes:** Correspondence should be directed to: WP, MT. These authors contributed equally to this work (Małgorzata Piechota, Katarzyna Banasiak, Nilesh Shanmugam).

## Abstract

Organismal functionality and reproduction depend on metabolic rewiring and balanced energy resources. However, the crosstalk between organismal homeostasis and fecundity, and the associated paracrine signaling mechanisms are still poorly understood. Using the *Caenorhabditis elegans* we discovered that large extracellular vesicles termed exophers, attributed in neurons and cardiomyocytes to the removal of damaged subcellular components, are released by body wall muscles to support embryonic growth. We found that exopher formation (exopheresis) is a non-cell autonomous process regulated by egg formation in the uterus. Our data suggest that exophers serve as transporters for muscle-generated yolk proteins used for nourishing and improving the growth rate of the next generation. We propose that the primary role of muscular exopheresis is to stimulate the reproductive capacity, thereby influencing the adaptation of worm populations to the current environmental conditions.

## Introduction

The proper cellular function relies on the removal of unwanted contents by proteolysis or degradation mainly via the ubiquitin-proteasome system (UPS) and autophagy (Dikic, 2017). Recently a complementary mechanism was described in *Caenorhabditis elegans*. Under proteotoxic stress, worms’ neurons can remove protein aggregates and damaged mitochondria via large membrane-surrounded vesicles called exophers. This process is beneficial for neurons functionality as cells that generate exophers perform better than those devoid of this mechanism (Melentijevic *et al*, 2017). However, not only neurons can remove cellular content via exophers. Murine cardiomyocytes can eject the subcellular content along with the mitochondria through exopher-like structures that are ultimately uptaken and eliminated by cardiac macrophages (Nicolás-Ávila *et al*, 2020). This extrusion phenomenon constitutes a significant but still poorly explored metabolic waste management pathway. Here, we report that *C. elegans* body wall muscle cells (BWM) can produce exophers in a sex-specific manner. Furthermore, the generation of muscular exophers depends on developing embryos in worms’ uterus and positively correlates with the number of retained eggs. Finally, we show that exophers serve as transporters for muscle-generated yolk proteins, which support offspring development. We identified a new role for exopheresis, which goes beyond the removal of proteotoxic cellular components, and has a transgenerational effect on the animal population.

## Results and Discussion

### *C. elegans* muscles remove cellular content via exophers

Using worms expressing fluorescent reporters in the body wall muscle cells, we identified muscle-derived vesicles resembling exophers. These reporters allow simultaneous tracking of mitochondria, which can be extruded in neuronal and cardiac exophers (Melentijevic *et al.*, 2017) and proteasomes, a key component of the proteostasis network. Similar to *C. elegans* neurons, muscles generate a small fraction (approximately 12%) of the vesicles that contain the mitochondria, but all of those we examined were filled with proteasomal subunits that are components of the 19S and 20S proteasome (**Figure 1A**, **S1A**). The size of the described vesicles extruded by BWM also corresponds to the neural exophers (Melentijevic *et al.*, 2017). However, those of muscular origin are generally more abundant and prominent (with diameters ranging from 2 to 15 μm) (**Figure 1B**). As with neural, exopheresis in muscles is not limited to a particular reporter or transgenic line (**Figures 1A-C, 2C, 4H**). Moreover, our electron microscopy (EM) studies visualized the membrane surrounded exophers adjacent to the BWM. Vesicles analyzed by EM show similar compactness, with the characteristic presence of mitochondria (**Figure 1C, S1C-D**). Most of them showed marked morphology changes, characterized by increased area and disturbed cristae organization as in cardiac exophers. However, we noted the presence of probably not compromised mitochondria in BWM-derived structures, suggesting that they are not only used to transport defective organelles. To finally confirm that the identified vesicles are the counterpart of exophers, we tracked their formation in response to RNAi knockdown of genes that regulate neuronal exopheresis (Melentijevic et al., 2017). Indeed, the appearance of muscle-derived exophers was also affected by depletion of EMB-8 cytochrome reductase and actin-binding protein POD-1, both linked to the generation of cell polarity (**Figure S1B**).

**Figure 1.**
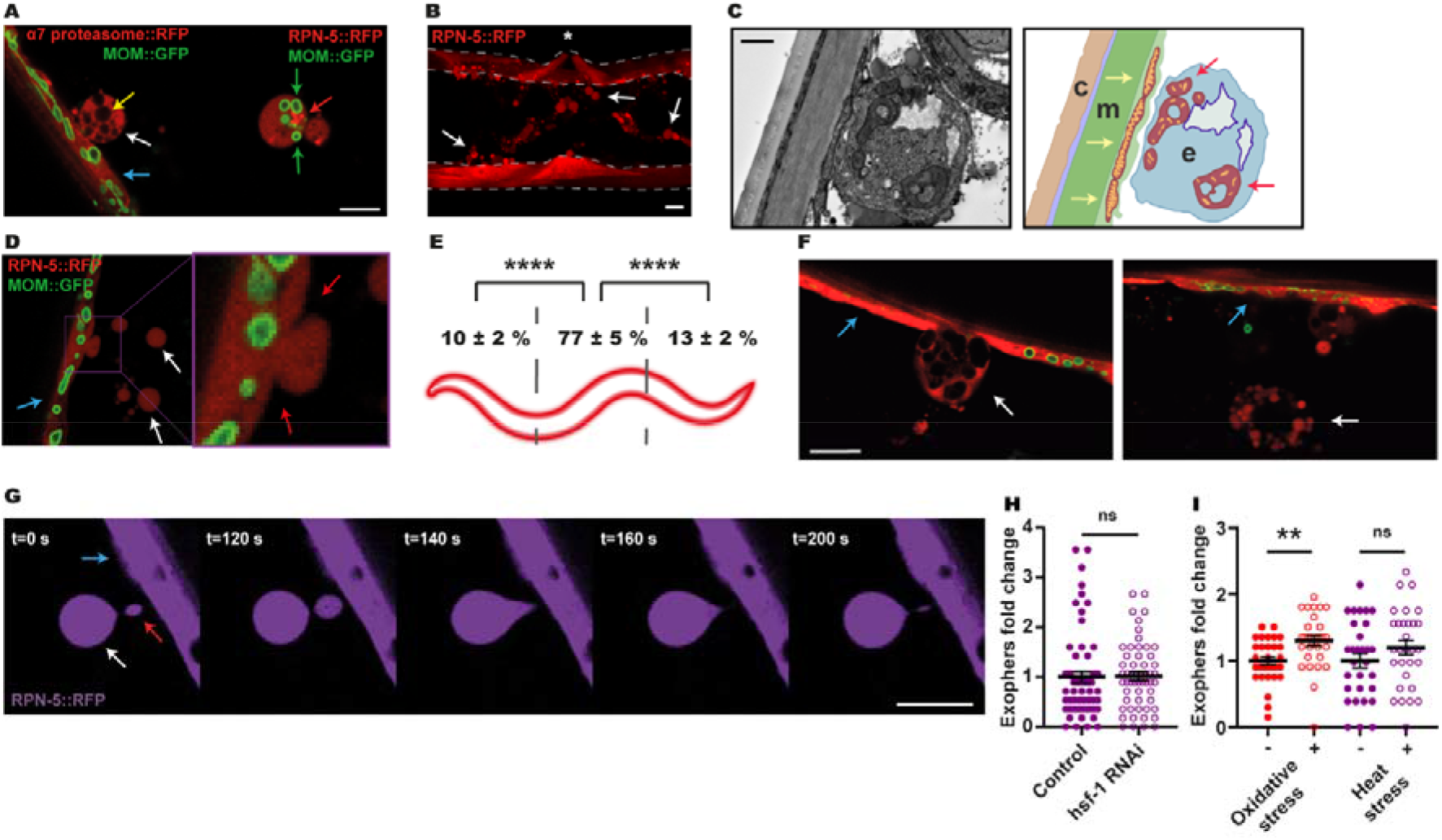
*C. elegans* muscles expel cellular content via exophers. (A) Muscular exophers contain organelles and large protein complexes. Arrows: white – exopher, blue – muscle cell, green – mitochondria, red – proteasome foci, yellow – unidentified vesicle. MOM – mitochondrial outer membrane. (B) BMW actively releases significant amounts of exopher. The image shows the middle part of the worm’s body with muscles marked with dashed lines. Arrows indicate representative exophers, and the asterisk indicates the position of the vulva. (C) The ultrastructure of the muscular exopher and its schematic view. Arrows: red – morphologically changed mitochondria inside the exopher, yellow – normal, elongated mitochondria inside muscle cell. c – cuticle, m – muscle, e – exopher. (D) Exophers are formed via a pinching-off mechanism. Arrows: white – exopher, blue – muscle cell, red – distorted muscle cell membrane during exopher formation. (E) Production of muscular exophers is not evenly distributed across all muscle cells. The highest number of exophers is produced by the muscles adjacent to the vulva. n = 46 animals; three biological replicates. (F) Large exophers may spontaneously disintegrate into smaller exophers. Arrows: white – exopher, blue – muscle cell. (G) Exophers may remain connected to the sending BWM cells via thin elastic tubes that allow further transfer of cellular material. Arrows: white – exopher, blue – muscle cell, red – cellular material transferred to exopher via elastic tube. (H) Proteostasis disruption by *hsf-1* knockdown does not increase exopher production. n = 60 and 55 animals, two biological replicates. (I) Proteostasis induced via oxidative stress but not via heat stress increases exopher production. n = 30 animals; three biological replicates. Scale bars are 10 μm (A, B, F, G) and 1 μm (C). Data are shown as mean ± SEM. ns - not significant, ** P < 0.01, **** P < 0.0001; (E, H, I) Mann-Whitney test.

**Figure 2.**
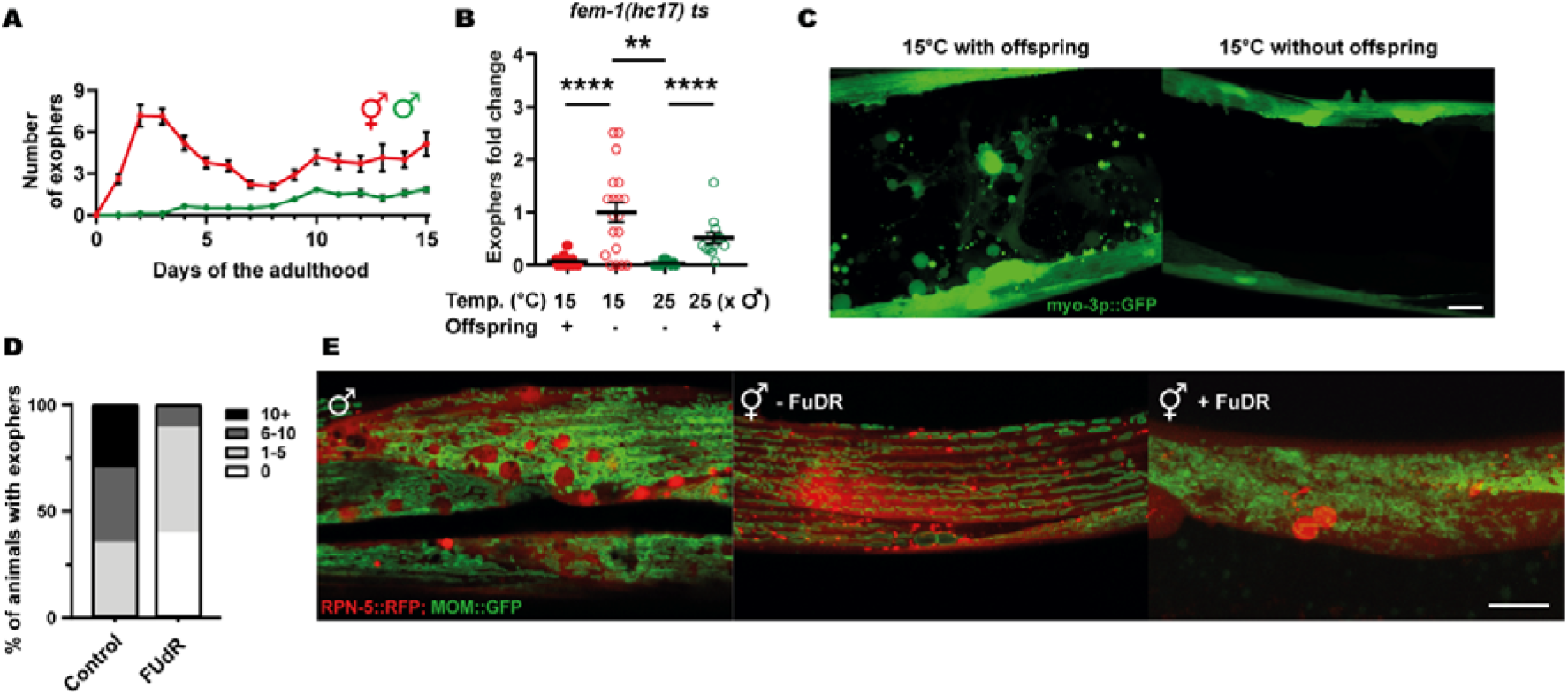
Exopher formation is sex-specific and fertility-dependent. (A) The highest number of exophers is produced during the hermaphrodite reproductive period and in aging animals. Males do not produce exophers during the first days of adulthood and begin to generate a small number of exophers later in life. Starting n = 90 hermaphrodites and 150 males; three biological replicates. (B) Feminized hermaphrodites of thermosensitive a *fem-1* mutant strain do not produce exophers regardless of growth temperature. This phenotype can be partially rescued by mating *fem-1* mutants with males. n = 10 - 26 animals; one biological replicate. (C) Repres entative images of the middle part of the worm body in panel B. (D) Hermaphrodites sterilized via FUdR treatment produce no exophers or only a few per animal. n = 17 and 20 animals; two biological replicates. (E) Males and sterile hermaphrodites (via FUdR treatment) show the formation of spherical structures in the BWM that resemble mature exophers. MOM – mitochondrial outer membrane. Scale bars are 10 μm. Data are shown as mean ± SEM; ** P < 0.01, **** P < 0.0001; (B) Mann-Whitney test.

Exophers are formed in the muscle cell and expelled outside via a pinching off mechanism (**Figure 1D**), and the majority are generated by adult hermaphrodite mid-body muscles (**Figure 1E**). Some exophers that bud off remain connected with the extruding BWM via a thin but elastic tube that permits the continued transfer of a large amount of cellular material into the extruded vesicle (**Figure 1G, Video S1**), similar to neuronal exophers (Melentijevic *et al.,* 2017). We also observed that large exophers could fragment into smaller vesicles (**Figure 1F**). Proteostasis impairment significantly increases neuronal exopher output (Hualin *et al,* 2019; Melentijevic *et al.,* 2017). In contrast, the number of muscular exophers did not change in response to depletion of the central proteostasis transcription factor HSF-1 (via *hsf-1* RNAi) or heat stress, and the number of exophers increased slightly under conditions of oxidative stress (**Figure 1H-I**). These observations suggest that proteostasis regulation might not be the core function of muscle exopheresis.

### Muscular exopheresis is a sex-specific process regulated in a non-cell autonomous manner

Next, we assessed the number of exophers at different time points of the *C. elegans* hermaphrodite life cycle. Reminiscent of neuronal exophers, muscular exophers are not produced during the larval stages, and their maximum level is reached around the second and third days of hermaphrodite adulthood (**Figure 2A**). Because this time point coincides with the worm’s maximum reproductive rate, we wondered if reproduction could influence exopher formation. To examine this possibility, we followed exopheresis in males. For the first three days of adulthood, males did not produce any exophers (**Figure 2A**). This finding suggests that germ cell maturation in the reproductive system of hermaphrodite worms, the process of oocyte fertilization, or embryonic development might regulate muscle exopheresis. To test these hypotheses, we took advantage of a thermosensitive *C. elegans fem-1* mutant strain that does not produce viable sperm at the restrictive temperature of 25 °C. At the permissive temperature of 15 °C, some animals can reproduce like wild type hermaphrodites, whereas the rest of the population is sterile (Nelson *et al,* 1978). The offspring-producing *fem-1* mutant animals grown at 15 °C generated a high number of muscular exophers. Contrarily, animals raised either at 15 °C or 25 °C that were unable to fertilize oocytes did not activate muscular exopheresis (**Figure 2B-C**), indicating that neither the female gonad nor the temperature itself was sufficient to trigger exopher release. However, re-establishing fertility of *fem-1* mutants by mating them to *him-*5 mutant males restored exopher production at the restrictive temperature (**Figure 2B**). Moreover, hermaphrodites sterilized via fluorodeoxyuridine (FUdR) treatment (Hosono, 1978) extruded no exophers or only a few per animal (**Figure 2D**), suggesting that the occurrence of developing embryos could indeed stimulate muscular exopheresis. We also found that FUdR-treated hermaphrodites often contained exopher-like structures in their BWM (**Figure 2E**, middle and right panels). Interestingly, we detected objects resembling non-extruded exophers in the BWM of males (**Figure 2E**, left panel), indicating that males are devoid of mechanisms triggering their expulsion.

The above results suggest that the occurrence of developing embryos could induce muscular exopheresis. Consistently, we observed a positive correlation between the number of exophers released and the number of eggs present in the worm uterus (**Figure 3A**). To further explore this link, we depleted the mRNA of genes responsible for various processes associated with egg-laying. RNAi depletion of the G-protein signalling gene *goa-1* leads to hyperactive egg-laying behaviour, resulting in the presence of fewer early-stage eggs within the uterus (Bany *et al*, 2003). As expected, *goa-1* knockdown caused a significant drop in the level of exopher release. In contrast, the egg-laying defects induced by *egl-1* and *egl-4* RNAi, which lead to egg retention in the uterus (Hirose *et al*, 2003), increased exopher formation by muscle cells (**Figure 3B**). In the absence of food, worms halt egg-laying and retain fertilized eggs in the uterus (Daniels *et al*, 2000; Dong *et al*, 2000); therefore, we anticipated that worms would generate more exophers when experiencing a food shortage. Indeed, the accumulation of developing eggs in the uterus caused by the transfer of adult worms to food-free plates resulted in a significant increase in muscle exopher secretion in contrast to L4 larvae grown under conditions of food shortage (**Figure 3C**).

**Figure 3.**
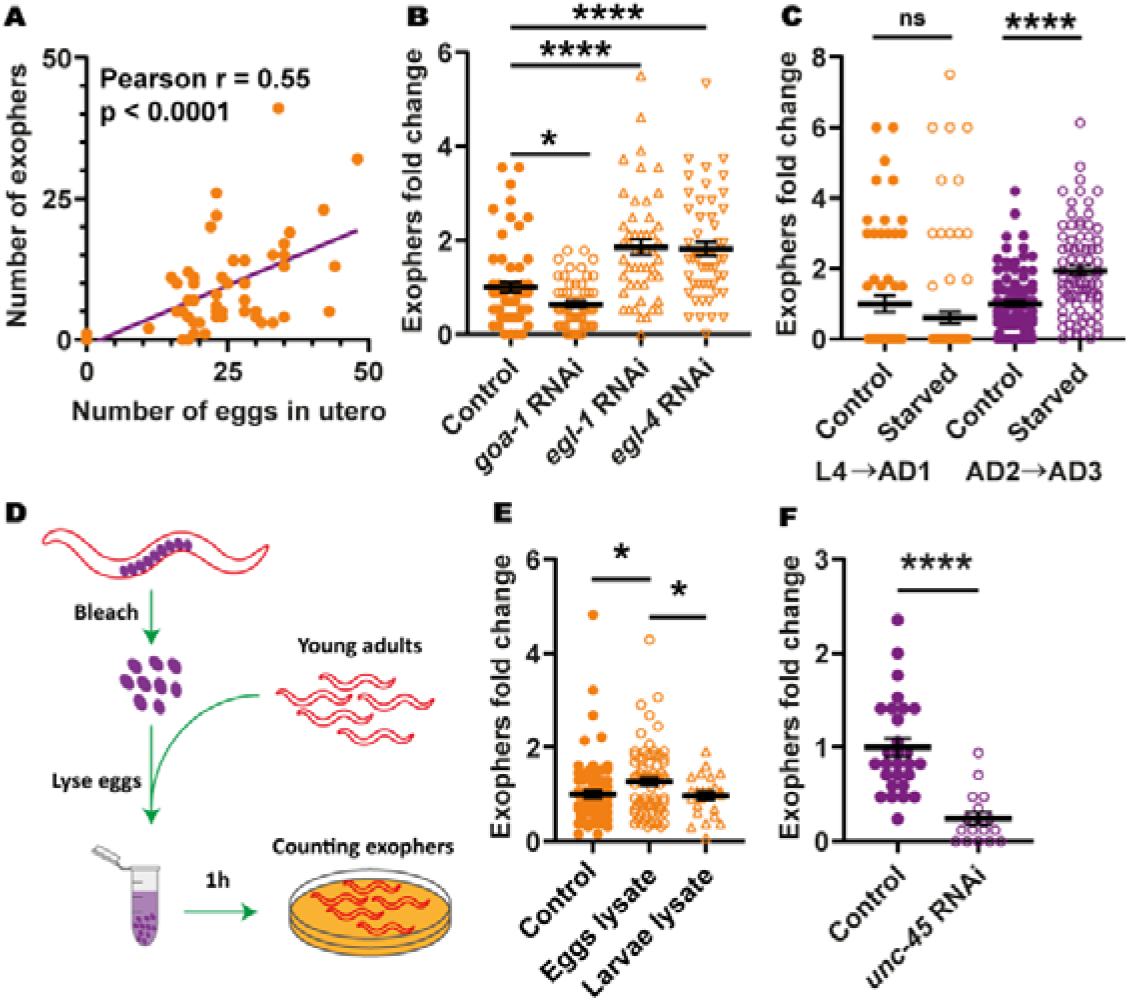
Muscular exopheresis is a non-cell autonomous process regulated by *in utero* developing embryos. (A) The number of produced exophers positively correlates with the number of *in utero* embryos. The violet line is a linear regression line, and each orange point represents one animal. All animals were 1 - 3 days old; n = 54 animals; three biological replicates. (B) RNAi knockdown of genes regulating the egg laying rate and their presence in the uterus influences exopher production. n = 50 - 60 animals; two biological replicates. (C) Eggs retention in the uterus caused by starvation during worm’s reproductive period increases exopher production. n = 81 - 90 animals; three biological replicates. (D) Schematic representation of the experimental setup for investigating the influence of egg lysate on exopher production. (E) Exposure of young-adult worms to embryo-derived substances increases exopher production. n = 24 - 77 animals; two biological replicates. (F) RNAi knockdown of the essential muscular co-chaperone *unc-45* reduces exopher production. n = 28 and 17 animals; two biological replicates. Data are shown as mean ± SEM; ns - not significant, * P < 0.05, **** P < 0.0001; (B, C, F) Mann-Whitney test; (E) two-tailed Welch’s t-test.

Next, we tested whether worm embryos could directly induce exopher production. We exposed young adult worms to extract from developing eggs or L2/L3 larvae of the wild-type strain (**Figure 3D**). Intriguingly, worms placed in contact with material derived only from lysed embryos increased exopheresis significantly (**Figure 3E**). This observation suggests that molecules that diffuse from embryos *in utero* are responsible for exopheresis induction. Finally, we decided to confirm that even in the face of disturbed proteostasis specifically in the BWM, a signal associated with the developing embryos in the uterus would be the primary regulator of muscular exopheresis. To this end, we knocked down the myosin-directed chaperone UNC-45, which results in severe defects in the sarcomeric proteostasis, as well as embryonic defects in cytokinesis and polarity determination (Gazda *et al*, 2013; Kachur *et al*, 2004; Pokrzywa & Hoppe, 2013). We initiated UNC-45 depletion in L4 larvae, which first leads to a disturbance of proteostasis in BWM, and later, in young gravid adults, inhibiting embryonic development. Despite the dysfunction of a myosin-chaperone network in muscle cells, which was indicated by complete paralysis of worms, we observed a dramatic inhibition of exopher formation (**Figure 3F**), highlighting the predominant role of maturating eggs in muscular exopheresis.

### Exophers transport muscle-synthesized yolk proteins and supports offspring development

The high number of exophers produced during the *C. elegans* lifespan by BWM leads to removing a rather significant portion of subcellular components by a single tissue. Thus, we hypothesized that this process should have substantial effects on worm muscle functionality and healthspan. To this end, we selected three types of worms from the synchronized population of gravid adults based on the number of extruded exophers, i.e., few (< 2), many (> 20), and control (6-8) animals (**Figure 4A**) and analyzed their locomotion. In neurons, exophers production is correlated with improved cell functionality; however, worms with intensified exopheresis did not show enhanced mobility (**Figure 4B, S2**). On the contrary, these animals displayed a reduction in exploratory locomotion state (**Figure 4C-D**). Since the production of exopher does not seem to have a positive effect on the functionality of the muscles, we next wondered whether, in connection with the role of embryos in exopheresis, this process benefited the offspring. Previous work showed that neuronal exopher content could be transported through the worm body to reach distant scavenger cells (coelomocytes) (Melentijevic *et al.,* 2017). However, the fate of muscle exophers may be different from neuronal counterparts, as their processing is not dependent on the apoptotic engulfment pathway proteins CED-1 and CED-6, the depletion of which did not change the muscle exopheresis level (**Figure S3A**). Muscle-specific transcriptomic analysis revealed the presence of significant levels of vitellogenin mRNAs (i.e., *vit-2, −5* and *-6)* (Blazie *et al,* 2015). Therefore, we hypothesized that the yolk components from BWM are transported through exophers to be used as a source of raw materials for maturating eggs. To address this possibility, we used RNAi knockdown of *vit-1* (a vitellogenin-coding gene) to reduce the level of the principal intestinal yolk protein (Perez & Lehner, 2019) and investigated whether it led to an increase in exopher biogenesis as a possible compensatory mechanism. Indeed, the number of accumulated muscle-released exophers nearly doubled in response to yolk protein depletion (**Figure 4E**). Moreover, the intensification of exopheresis in the mother increased the amount of vitellogenin in developing eggs (**Figure 4F**). In contrast, depletion of the RME-2 yolk receptor and subsequent inhibition of yolk uptake by oocytes led to a drastic reduction in exopheresis (**Figure 4G**), which cannot be explained by the sterility of worms, as the embryos’ viability is not entirely compromised even in the *rme-2* deletion mutant (b1008) (more than 20% of embryos are alive) (Grant & Hirsh, 1999). Next, we followed the localization of endogenous vitellogenin-2 (VIT-2) fused to GFP (CRISPR/Cas9 knock-in strain) and muscle exopher markers in the hermaphrodite. We detected the presence of endogenous VIT-2::GFP in the BWM of day-2 adult worms, as well as a significant accumulation in many muscle exophers (**Figure 4H, S3B**). These results suggest that exophers can mediate the transport of additional portions of muscle-produced vitellogenin, which is ultimately deposited in oocytes from the body cavity (Hall *et al*, 1999). Finally, we followed the growth of the progeny of hermaphrodites exhibiting different levels of exopheresis (**Figure 4A**). We found that offspring from mothers with a high number of muscle-released exophers grew faster (**Figure 4I**). This is in line with previous reports showing that yolk-reach eggs support animals post-embryonic survival and larvae development (Perez *et al*, 2017; Perez & Lehner, 2019; Van Rompay *et al*, 2015).

**Figure 4.**
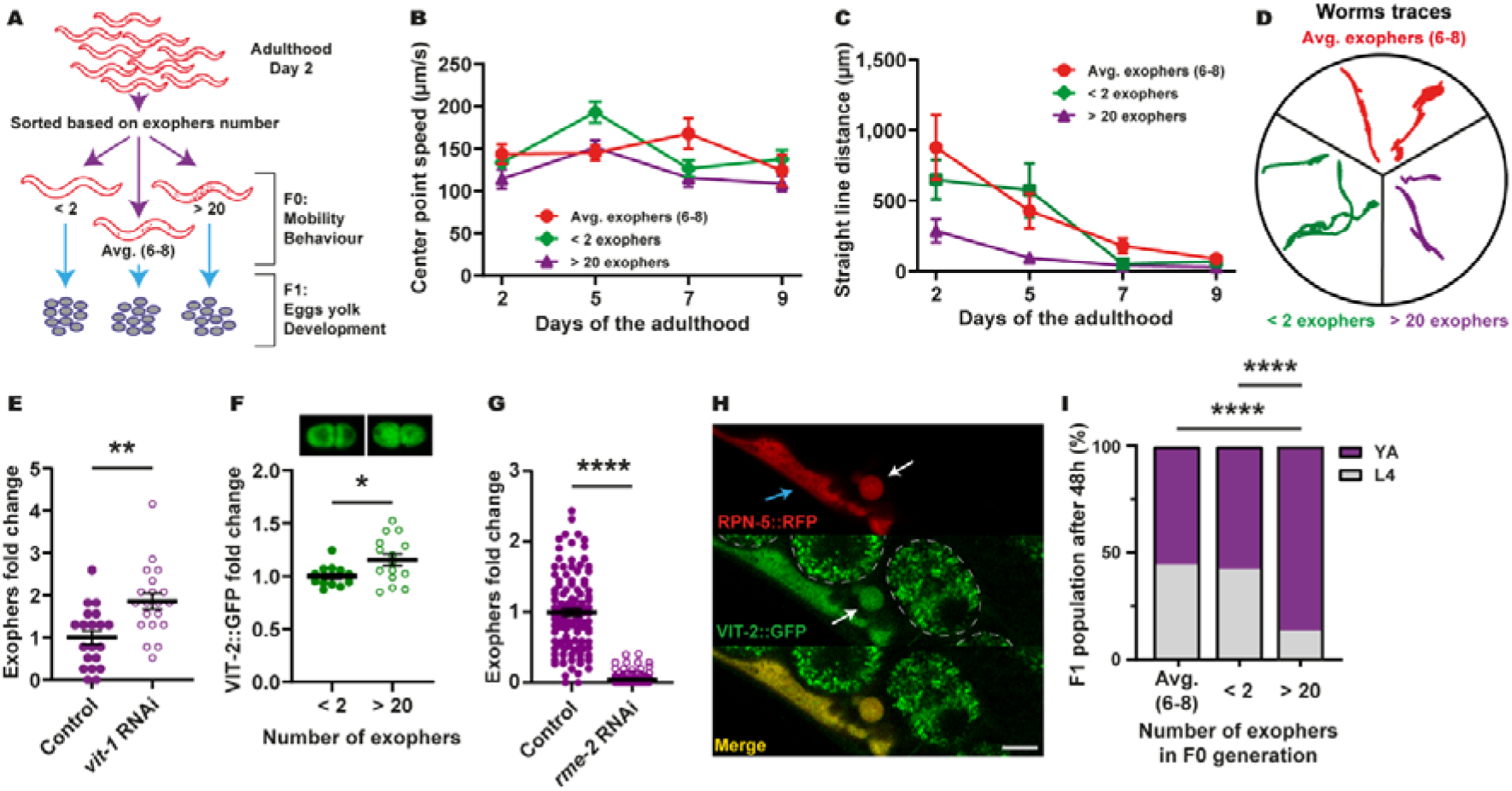
Muscular exopheresis benefits offspring development. (A) Schematic representation of the experimental setup for investigating the influence of overactive exopheresis on F0 and F1 worm generation. (B) Exopher production does not improve muscle functionality. n = 11 - 38; three biological replicates. (C-D) Animals with overactive exopheresis have reduced exploratory behaviour presented as a reduction in straight line distance travelled (C) and by representative worms traces on the plate (D). n = 11 - 34; three biological replicates. (E) RNAi knockdown of the egg yolk precursor protein VIT-1 increases the number of muscular exophers. n = 20 animals; two biological replicates. (F) Embryos from hermaphrodite mothers that produce a high number of exophers contain more egg yolk precursor protein VIT-2. Representative images of embryos with endogenous VIT-2::GFP levels from mothers with different exopheresis activity are shown above the graph. n = 15 and 14 eggs, five and six biological replicates (animals). (G) RNAi knockdown of RME-2 yolk receptor abolish exopher production. n = 110 – 123 animals; three biological replicates. (H) Muscle-produced VIT-2 is released from muscles via exophers. The image shows the midbody of worms expressing the proteasome subunit RPN-5 tagged with RFP in BWM and VIT-2::GFP endogenous expression. Arrows: white – exopher, blue – muscle cell. Dashed lines mark eggs present in the uterus. Scale bar is 10 μm. (I) Offspring of worms with overactive exopheresis develop faster. YA – young adult stage, L4 – last larval stage. n = 317 – 372 animals; two biological replicates. Data are shown as mean ± SEM. * P < 0.05, ** P < 0.01, **** P < 0.0001; (E, G) Mann-Whitney test; (F) two-tailed Welch’s t-test; (I) Fisher’s exact test.

Exophers result from the activation of non-canonical mechanisms that allow the neuronal and muscle cells to release the sub-cellular components together with organelles such as mitochondria or nano-organelles such as proteasomes. Accordingly, based on their content and participation of the autophagy machinery in exophers formation, this vesicle can be considered part of the proteostasis network. Here we show that exophers are not only a storage compartment of cellular waste, but the muscular exopheresis in *C. elegans* represents a previously uncharacterized nutrient management program associated with nourishing the next generation of worms. Activation of exopheresis occurs with the first appearance of developing eggs in the uterus and intensifies in environmental food depletion situations. Hermaphrodites utilize a narrow time window between 2-4 days of adulthood to simultaneously expel probably parts of the dysfunctional cell elements out of the muscles, as well as material that can be used by the embryos. Likewise, the disturbance of yolk synthesis increases exopher biosynthesis. Consequently, in mothers with highly active exopheresis, the volume of yolk content in eggs increased. Our results show that yolk protein produced in BWM is transferred to exophers and ultimately delivered to oocytes. Therefore, the use of exophers for the transport of vitellogenin represents an elegant mechanism by which remote cells can enrich the nourishment of developing embryos. This process leads to the production of larvae better prepared to thrive in the current environmental conditions. Thus, muscular exopheresis is likely an adaptive mechanism that affects the dynamics of population growth. The impact of exopheresis on early reproduction may be particularly crucial for wild worms, given the significantly shortened life expectancy observed under more natural conditions (Van Voorhies *et al.,* 2005). In addition, yolk protein can be synthesized in the muscles of oviparous animals like zebrafish (Zhong *et al,* 2014), and is supplemented from the mother to the intraovarian embryo in viviparous animals (Iida *et al,* 2019). Hence, it is tempting to speculate that the role of muscular exopheresis in supporting progeny development could be evolutionarily conserved. However, the exact mechanisms by which oocyte fertilization and subsequent embryonic development initiate exopher formation and how this exopheresis is executed at the molecular level require further studies.

---

**Figure S1.**
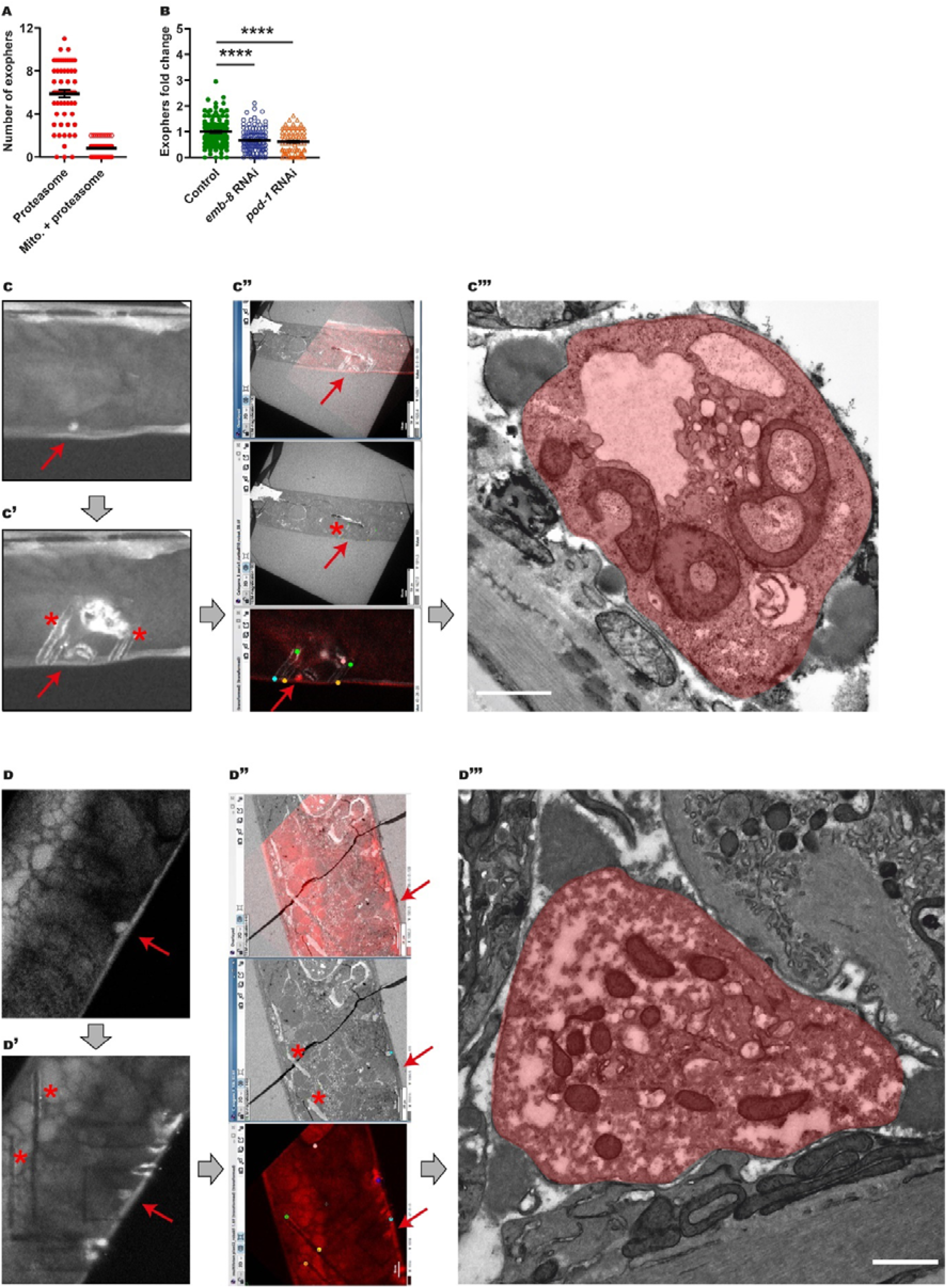
Ultrastructure of muscle exophers, the formation of which depends on cell polarity regulating proteins. (A) Small fraction of muscular exophers contain mitochondria. n = 60; six biological replicates. (B) The formation of muscle-derived exophers is affected by depletion of proteins linked to the generation of cell polarity. n = 80 - 120; *pod-1* - two biological replicates, *emb-8* RNAi - three biological replicates. Data are shown as mean ± SEM. **** P < 0.0001, Mann-Whitney test. (C-D) Exophers visualization using correlative light electron microscopy (CLEM). Exopher’s red fluorescence (red arrow). (C’-D’) Exopher (red arrow) surrounded by marks (asterisk) after NIRB, autofluorescence of branded frame edges is visible. (C”-D”) CLEM tracking the exopher - LM image, EM image and their overlay. (C”‘-D”‘) Image shows exopher’s (coloured in red) ultrastructure. Thick, grey arrows show order of steps done for the final exopher detection under the electron microscope. Scale bars are 1 μm.

**Figure S2.**
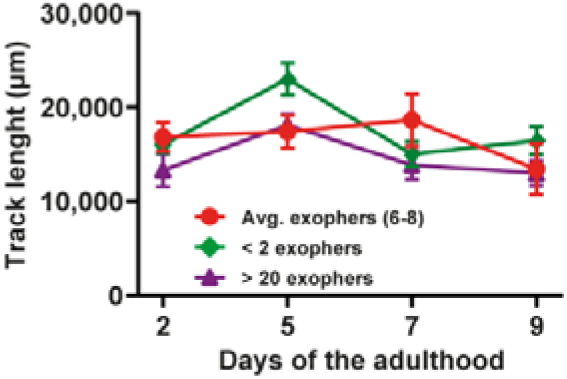
Active exophoresis affects worm mobility. Graph shows the track lengths of animals exhibiting different exopheresis activities. n = 11 - 38; three biological replicates. Data are shown as mean ± SEM.

**Figure S3.**
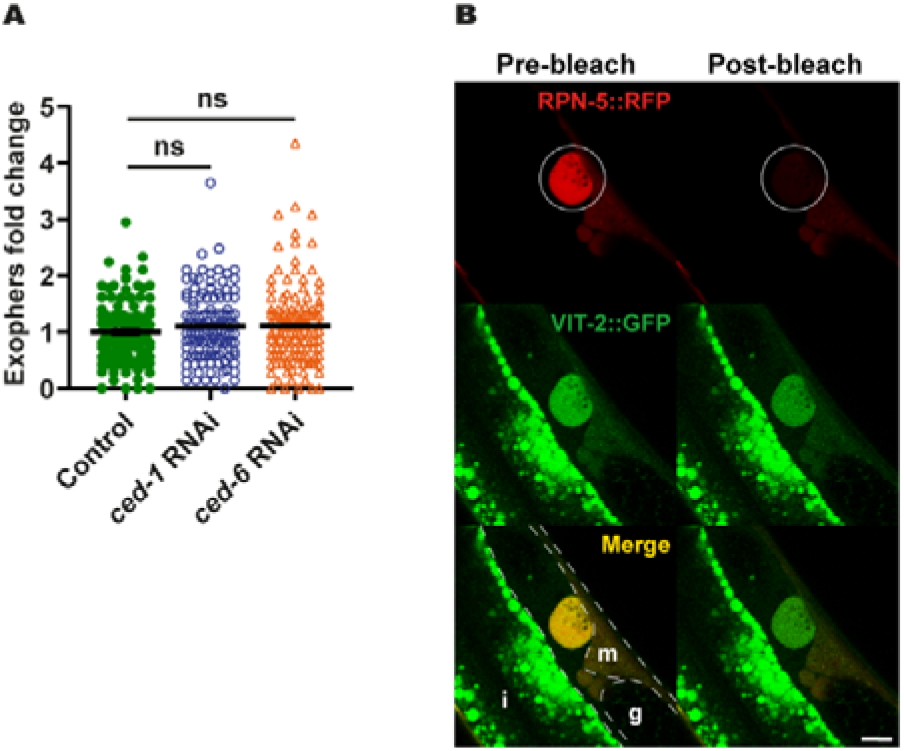
Exophers contain muscle-produced vitellogenin. (A) Muscular exophers formation does not depend on the apoptotic engulfment pathway proteins CED-1 and CED-6. n = 114 - 120; three biological replicates. Data are shown as mean ± SEM. ns - not significant, Mann-Whitney test. (B) Images show the formation of an exopher filled with the proteasome and vitellogenin. Images were captured before and after RPN-5::RFP photobleaching, confirming that the high signal from endogenous VIT-2::GFP in forming exophers is not an imaging artefact. The white circle marks the area that was bleached using a 555 nm laser and the position of developing exopher. Dashed lines mark different tissue borders: m – muscle, i – intestine, g – gonad. Scale bar is 10 μm.

## Methods

### Data reporting

No statistical methods were used to predetermine the sample size. The experiments were not randomized. The investigators were not blinded to allocation during experiments and outcome assessment except for exploratory behaviour experiments.

### Worm maintenance and strains

Worms were maintained on Nematode Growing Medium (NGM) plates seeded with OP50 *Escherichia coli* bacteria at 20 °C unless otherwise stated (Brenner, 1974). A list of all strains used in the study, together with the information in which experiments they were used is provided in Supplementary Table 1.

### Generation of plasmids

All constructs were cloned using the SLiCE method (Zhang *et al,* 2012) and were sequenced for verification. Construct for the expression of GFP on the mitochondrial outer membrane in the worm’s body wall muscles was prepared as follows. First, destination vector pMT26 containing pCG150 vector backbone with the *myo-3* promoter and *unc-54* 3’UTR sequences separated by KspAI restriction site was made. Next, codon-optimized sequences (Redemann *et al,* 2011) for GFP, linker containing attB5 sequence, and the sequence of the first 55 amino acids from TOMM-20 protein (Watanabe *et al*, 2011) (from a plasmid that was synthesized for this study) were PCR amplified and inserted into linearized pMT26 vector. To generate construct for expression of RPN-5 or PAS-7 tagged with RFP in worm’s body wall muscles, sequences encoding respective proteins were PCR amplified and inserted into pMT26 linearized destination vector. As a template for the *rpn-5*-coding sequence, we used plasmid bearing codon-optimized *rpn-5* cDNA with 3 artificial introns, which was synthesized for this study. The sequence of *pas-7* gene was directly amplified from N2 worms genomic DNA, and the sequence of RFP (wrmScarlet) was amplified from pSEM89 plasmid (El Mouridi *et al,* 2017).

### Transgenic strains generation

Worms transgenic strains were created by microparticle bombardment using *unc-119(ed3)* rescue as a selection marker (Praitis *et al,* 2001). After phenotypic confirmation of successful plasmid insertion, transformants were backcrossed two times against N2 strain to remove the *unc-119(ed3)* background.

### Scoring exophers and fluorescence microscopy

For scoring of exophers, a confocal microscope or a stereomicroscope was used. When using the confocal microscope, animals were transferred onto 3 % agarose pads prepared in H_2_O and formed on a microscope slide. Next, animals were immobilized on the pad using 6 μL of PolySciences 0.05 μm polystyrene microspheres or 25 μM tetramisole and covered with a glass coverslip. Immediately afterward, animals were imaged using an inverted Zeiss LSM800 laser-scanning confocal microscope with 40x or 63x oil immersion objectives. The 488 nm and 561 nm lasers were used for excitation of GFP and RFP fluorescent proteins, respectively. Z-stacks, which covered the whole animal, were taken and the number of exophers released by muscles was counted. Finally, data were normalized to the average number of exophers present in control animals and compared between conditions.

For scoring exophers with the stereomicroscope, we used Leica M165FC stereomicroscope equipped with Leica EL6000 lamp and standard Texas Red and GFP filter sets. Age synchronized, freely moving, day-2 adult animals were directly visualized on culturing NGM plates and the number of visible exophers released by muscles were counted. Finally, data were normalized to the average number of exophers present in control animals and compared between conditions.

Representative pictures of exophers presented in the manuscript were taken with the usage of an inverted Zeiss 700 laser-scanning confocal microscope equipped with a 40x oil objective. The 488 nm and 555 nm lasers were used for excitation of GFP and RFP fluorescent proteins, respectively. To investigate the presence and distribution of exophers, z-stacks were taken and processed with ZEN software.

### Electron microscopy

#### EM sample preparation

*C. elegans* day 2 adult worms were fixed with 2% paraformaldehyde (Sigma Aldrich, P6148) and 1% glutaraldehyde (Sigma Aldrich, EM grade) in 0.2M Hepes pH 7.3 overnight. Next, the samples were washed three times in 0.2M Hepes pH 7.3, followed by overnight incubation with 0.1% ruthenium red solution in distilled water. After staining, worms were mounted with 2% gelatin on gridded 35mm dishes (MatTek 35mm dish, No. 1.5 Gridded Coverslip 14 mm Glass Diameter, P35G-1.5-14-CGRD) and designated for branding.

#### Near infra-red branding (NIRB)

Worms were localised on a gridded dish in bright field mode. Next, the exophers’ red fluorescence was imaged using multiphoton Examiner.Z1 LSM 7MP microscope equipped with LSM NDD detectors and water 20x NA 1.0 objective (Zeiss, Oberkochen, Germany). For the imaging, pulsed laser at 1020 nm (Coherent Chameleon) and 600-700 nm detection filter were used. Z-stacks of the samples were acquired using 4 times averaging in the line mode, pixel size of 138 x 138 nm and 500 nm interval in the Z axis. For each sample a Z-stack was acquired before and after the branding.

The ROI was marked by NIRB (Bishop *et al,* 2011). The frame contour was bleached below the place of interest with a pulsed laser at 910 nm (Coherent Chameleon, 2 W at 910 nm), using 20-40 iterations with 20% of the laser power and pixel dwell time of 6-25 μs. The final parameters (number of iteration and pixel dwell time) were adjusted separately for each sample. The branding was repeated until the edges of the frame were highly fluorescent at 910 nm.

#### EM processing

Branded samples were prepared for electron microscopy according to a published protocol(Deerinck *et al,* 2010) with minor changes (Śliwińska *et al,* 2020). Briefly, animals were post-fixed with 1% aqueous solution of osmium tetroxide (Polysciences Europe GmbH 0972B-5) and 1.5% potassium ferrocyanide (Sigma Aldrich, St. Louis, MO, USA, P3289) in phosphate buffer for 30 min on ice. Next, samples were immersed in 1% aqueous thiocarbohydrazide (Sigma Aldrich, St. Louis, MO, USA, #88535) for 40 min, post-fixed with 2% aqueous solution of osmium tetroxide for 60 min (all at room temperature) and incubated in 1% aqueous uranyl acetate at 4°C overnight. The next day, samples were immersed in 0.66% lead aspartate for 60 min at 60°C, dehydrated with increasing dilutions of ethanol, infiltrated with Durcupan resin (Sigma Aldrich, St. Louis, MO, USA, #44610), embedded using BEEM capsules according to a published protocol (Hanson *et al,* 2010) and cured at 70°C for 72 h.

The resin blocks were trimmed and cut with an ultramicrotome (ultracut R or EM UC7, Leica) and ultrathin sections (70 nm thick) were collected on formvar-coated copper grids, mesh 100 (Agar Scientific, AGS138-1 or EMS FF100-CU-50) or copper slot grids (EMS FCF 2010-CU-SB-50). For serial block face scanning electron microscopy trimmed pyramids were cut off with a razor blade, mounted to aluminium pins (Gatan system pins, Micro to Nano, Netherlands, 10-006003-50) with cyanoacrylate glue, trimmed, block face polished and grounded with silver paint (Ted Pella, Redding, USA, 16 062-15).

#### Transmission Electron Microscopy Imaging

Specimen grids were examined with a transmission electron microscope JEM 1400 (JEOL Co., Tokyo, Japan, 2008), equipped with a 11 megapixel TEM camera MORADA G2 (EMSIS GmbH, Münster, Germany) at Nencki Institute of Experimental Biology PAS or with a Tecnai T12 BioTwin electron microscope (FEI, Hillsboro, OR, USA) equipped with a 16 megapixel TemCam-F416 (R) camera (TVIPS GmbH) at International Institute of Molecular and Cell Biology in Warsaw.

#### Exopher tracking

Identification of branded expoher was performed with eC-CLEM open source software (Paul-Gilloteaux *et al*, 2017). The NIRB frame is visible on both FM and EM image. The marks on EM image are placed on the frame’s edges and respectively its position is adjusted on FM image allowing for correlation of both images. These coordinates allow to set out exopher position on the TEM image.

### RNA interference

RNA interference in *C. elegans* was performed using the standard RNAi feeding method and RNAi clone (Kamath & Ahringer, 2003). For experiments, we used NGM plates supplemented with 1 mM IPTG and 25 μg/μl carbenicillin seeded with HT115 *E. coli* bacteria expressing double-stranded RNA (dsRNA) against the gene of interest or, as a control, with bacteria without a vector. Worms were placed on freshly prepared RNAi plates, either as age-synchronized pretzel-stage eggs, L1 larvae, or L4 larvae. The number of exophers was counted on day 2 adult worms using a confocal microscope or a stereomicroscope.

### Stress influence on exophers production

Worms were age-synchronized using alkaline hypochlorite solution (bleaching procedure), as previously described (Porta-de-la-Riva *et al*, 2012). The harvested eggs were incubated overnight at 16 °C for hatching. Approx. 1000 L1 larvae were transferred on NGM plates and incubated at 20 °C till they reached day-2 of adulthood. The worms were further channelled to the respective stress treatments.

#### Oxidative stress

Approx. 100, day-2 adult worms were washed-off from the NGM plates and rinsed 3 times with M9 buffer. The worms to be stressed were suspended in 1 ml of 5 mM hydrogen peroxide solution prepared in M9 buffer, whereas control worms were suspended in M9 buffer. The tubes were incubated on a shaker at 20 °C for 60 minutes.

#### Heat stress

Similarly, approx. 100, day-2 adult worms were washed-off form NGM plates and rinsed 3 times with M9 buffer. The worms were further suspended in a 1 ml M9 buffer. The worms to be heat stressed were incubated in a shaker at 33 °C for 60 minutes, whereas the control animals were incubated at 20 °C for 60 minutes.

#### Exophers quantification

From each stress/control treatment, 30 worms were picked onto agarose pad slides individually for the quantification of exophers. Exophers were quantified using a confocal microscope. Obtained data were normalized to the average number of exophers present in control animals and compared between conditions.

### Number of exophers on consecutive days

For each biological replicate, 30 L4 larvae hermaphrodites or 50 L4 larvae males were transferred to fresh NGM plates (5 hermaphrodites or 10 males per plate). For the next consecutive 15 days, the number of exophers in each animal was counted using a stereomicroscope. Hermaphrodites were transferred to fresh plates every 2 - 3 days. Males were kept on the same plate until the end of the experiment. All animals that died during the experiment time course were removed from the plate.

### Measuring exophers in *fem-1* mutant

Approx. 50 L1 larvae from *fem-1(hc17)ts* mutant strain expressing GFP in BWM were transferred to 2 fresh NGM plates. During the time course of the experiment, one of the plates was kept at 15 °C while the second one was kept at 25 °C. When worms reached L4 larval stage, each worm was transferred to a separate plate and grown at the same temperature as before. After 48 hours at 25 °C or 72 hours at 15 °C, using a stereomicroscope, a number of exophers released from worms muscles was counted and plates were scored for F1 offspring to assign all worms as fertile or infertile.

Additionally, a phenotype rescue experiment was performed for sterile animals grown at 25 °C. 24 hours after they reached the adulthood, single animals were placed on separate plates and mated with 6-8 *him-5* mutant day-1 males. After next 48 hours, a number of exophers released from worms muscles was counted.

### FUdR assay

Age-synchronized animals were placed on NGM plates seeded with OP50 *E. coli* bacteria as a food source until they reached young adulthood (day 0). Next, animals were selected and moved to test plates containing 25 μM fluorodeoxyuridine (FUdR) to prevent embryonic development and egg hatching (Mitchell *et al,* 1979) or control plates without FUdR. The number of exophers was counted on adult day-2 using confocal microscopy.

### Exophers and *in utero* eggs correlation

For correlating the number of exophers with the number of eggs present in worms uterus, 1 to 3-days old animals were used. First, muscular exophers for a single worm were counted using the stereomicroscope. Next, worm was transferred to a 10 μl drop of 1.8 % hypochlorite solution put on a microscope slide. Finally, after approx. 5 minutes when the hermaphrodite mother was fully bleached, the number of eggs released from worm’s uterus was counted. Data analysis was performed using GraphPad Prism 8 software.

### Starvation assay

To assess the influence of worms starvation on the exophers production, day-2 adult worms and L4 larvae were moved to bacteria-free NGM plates. After 24 hours of food deprivation, the number of exophers was counted using a stereomicroscope. As a control, day-3 adult worms and day-1 adult worms grown for the whole time on NGM plates seeded with bacteria were used. Obtained data were normalized to the average number of exophers present in control animals and compared between conditions.

### Eggs lysate assay

To obtain eggs, age-synchronized N2 gravid hermaphrodites were bleached using an alkaline hypochlorite solution. Harvested eggs were suspended in appropriate volume of M9 buffer to reach the concentration of approx. 200 eggs per μl. As a control, age-synchronized L2/L3 larvae were collected and suspended in M9 buffer to reach the concentration of approx. 25 larvae per μl. Next, eggs and larvae were flash-frozen in liquid nitrogen, thawed on ice, and sonicated 3 times for 10 seconds to obtain eggs and larvae lysate. 150 μl of eggs or larvae lysate was mixed with 150 μl of concentrated OP50 *E. coli* bacteria and placed in a 0.5 ml Eppendorf tube. Approx. 30 day-1 young adult hermaphrodites were transferred to Eppendorf tube containing a mixture of eggs or larvae lysate and bacteria. Next, the Eppendorf tube was placed on the rotator for 1 hour at room temperature. After 1 hour, the content of the tube was placed on the fresh NGM plate seeded with OP50 bacteria. 18 hours later the number of exophers in each worm was counted. For mock control animals, the whole protocol was the same except that for bleaching step no worms were used.

### Worm exploratory behaviour

Age-synchronized, day-2 adult worms which had an average (6-8), few (< 2), or many (> 20) exophers were sorted to separate NGM plates. Approximately 10 worms per replicate were brought onto NGM plates and the worm movement was recorded for 2 minutes using the WormLab system (MBF Bioscience). The frame rate, exposure time, and gain were set to 7.5 frames per second, 0.0031 s, and 1, respectively. The track length, straight-line distance, center point speed, and the overall track pattern of individual worms were analyzed using the WormLab software (MBF Bioscience).

### Vitellogenin levels in embryos

Vitellogenin levels in embryos were measured based on the GFP signal from fluorescently tagged VIT-2 protein. Approx. 200 late L4 larvae were transferred to fresh NGM plate. On day 2 of worms adulthood, the number of exophers in each animal was counted and worms with less than 2 or more than 20 visible exophers were transferred to new plates. Next, animals from each group were individually transferred to a 10 μl drop of M9 buffer which was placed on a microscope slide. Using a sharp injection needle, worm was cut open to release eggs from the uterus. Fluorescent signal from 2-cell embryo stage was captured using Leica M165FC stereomicroscope equipped with Leica EL6000 lamp, standard GFP filter set, and Leica DFC365 FX CCD camera. Magnification used for recording pictures was set to 192x. Exposure time and gain were set to 600 ms and 2, respectively. The fluorescent signal was quantified using Leica Las X software and was normalized to the average signal from eggs, which were obtained from animals with less than 2 exophers.

### Worms development assay

25 - 30 age-synchronized, day-2 adult worms which had an average (6-8), few (< 2), or many (> 20) exophers were sorted to separate NGM plates. Gravid adults were allowed to lay eggs for 4 hours, afterward removed from the plates, and the development of their offspring was followed. 46 hours later, using the stereomicroscope, developmental stage of each animal was checked, and the proportion between L4 larvae stage worms and young adult worms was calculated.

### Statistical analysis

Statistical tests used in this study were two-tailed Welch’s t-test and Fisher’s exact test. P-value < 0.05 was considered significant.

## Acknowledgements

We thank the Caenorhabditis Genetics Center (funded by the NIH National Center for Research Resources, P40 OD010440) for strains; T. Hoppe for expert advice and Carl Kutzner for RNAi clones; T. Wegierski for confocal microscopy assistance; B. Uszczyńska-Ratajczak, K. Szczepanowska, H. Bringmann, and members of Chacińska and Pokrzywa laboratories for discussions and comments on the manuscript. Multiphoton microscopy imaging, near infrared branding, and serial block face scanning electron microscopy imaging were performed at the Laboratory of Imaging Tissue Structure and Function at Nencki Institute of Experimental Biology PAS. Transmission electron microscopy imaging and ultramicrotomy was performed either at Core Facility of International Institute of Molecular and Cell Biology in Warsaw or using the equipment of Laboratory of Electron Microscopy at Nencki Institute of Experimental Biology PAS.

## Funding

Work in the W.P. laboratory was supported by the Foundation for Polish Science co-financed by the European Union under the European Regional Development Fund (grant POIR.04.04.00-005EAB/18-00 to K.B., K.K. and W.P.), the European Molecular Biology Organization (EMBO Installation Grant No. 3916 to M.P. and W.P.), the Polish National Science Center (grant UMO-2016/23/B/NZ3/00753 to N.S. and W.P.), and the Deutsche Forschungsgemeinschaft (DFG FOR 2743 to W.P.). Work in the A.C. laboratory was funded by “Regenerative Mechanisms for Health” project MAB/2017/2 (M.T. and A.C.) carried out within the International Research Agendas programme of the Foundation for Polish Science co-financed by the European Union under the European Regional Development Fund and was supported by a POLONEZ Fellowship of National Science Centre, Poland, 2016/21/P/NZ3/03891 (M.T.), within European Union’s Horizon 2020 research and innovation programme under the Marie Skłodowska-Curie grant agreement no. 665778.

## Author contributions

M.T., W.P., M.P., K.B., N.S., M.M., M.A.Ś., M.N., K.K. and N.N. conducted and designed experiments, M.T. and W.P. conceived the project and supervised the study. W.P., A.C. and M.T. secured the funding. W.P. and M.T. wrote the manuscript with input from A.C. and M.P.

## Author Information

The authors declare no competing financial interests. Further information and requests for resources and reagents should be directed to and will be fulfilled by the Lead Contact, Wojciech Pokrzywa.

## Supplementary Information

Supplementary Table 1

List of *C. elegans* strains used in this study

Supplementary Video 1

The video shows the cellular content being transfer from BWM cell to exopher via elastic tube.

## Notes

### Competing Interest Statement

The authors have declared no competing interest.

### Summary of Updates

- Figure 1-4, Supp. Fig. 1 and 3 revised and updated - Authors list and affiliations updated - Text revised

## References

Bany IA, Dong MQ, Koelle MR (2003) Genetic and cellular basis for acetylcholine inhibition of Caenorhabditis elegans egg-laying behavior. J Neurosci 23: 8060–8069

Bishop D, Nikić I, Brinkoetter M, Knecht S, Potz S, Kerschensteiner M, Misgeld T (2011) Near-infrared branding efficiently correlates light and electron microscopy. Nat Methods 8: 568–570

Blazie SM, Babb C, Wilky H, Rawls A, Park JG, Mangone M (2015) Comparative RNA-Seq analysis reveals pervasive tissue-specific alternative polyadenylation in Caenorhabditis elegans intestine and muscles. BMC Biol 13: 4

Brenner S (1974) The genetics of Caenorhabditis elegans. Genetics 77: 71–94

Daniels SA, Ailion M, Thomas JH, Sengupta P (2000) egl-4 acts through a transforming growth factor-beta/SMAD pathway in Caenorhabditis elegans to regulate multiple neuronal circuits in response to sensory cues. Genetics 156: 123–141

Deerinck TJ, Bushong EA, Thor A, Ellisman MH, 2010. NCMIR methods for 3D EM: A new protocol for preparation of biological specimens for serial block face scanning electron microscopy. National Center for Microscopy and Imaging Research, University of California San Diego, La Jolla, CA92093.

Dikic I (2017) Proteasomal and Autophagic Degradation Systems. Annu Rev Biochem 86: 193–224

Dong MQ, Chase D, Patikoglou GA, Koelle MR (2000) Multiple RGS proteins alter neural G protein signaling to allow C. elegans to rapidly change behavior when fed. Genes Dev 14: 2003–2014

El Mouridi S, Lecroisey C, Tardy P, Mercier M, Leclercq-Blondel A, Zariohi N, Boulin T (2017) Reliable CRISPR/Cas9 Genome Engineering in. G3 (Bethesda) 7: 1429–1437

Gazda L, Pokrzywa W, Hellerschmied D, Lowe T, Forne I, Mueller-Planitz F, Hoppe T, Clausen T (2013) The myosin chaperone UNC-45 is organized in tandem modules to support myofilament formation in C. elegans. Cell 152: 183–195

Grant B, Hirsh D (1999) Receptor-mediated endocytosis in the Caenorhabditis elegans oocyte. Mol Biol Cell 10: 4311–4326

Hall DH, Winfrey VP, Blaeuer G, Hoffman LH, Furuta T, Rose KL, Hobert O, Greenstein D (1999) Ultrastructural features of the adult hermaphrodite gonad of Caenorhabditis elegans: relations between the germ line and soma. Dev Biol 212: 101–123

Hanson HH, Reilly JE, Lee R, Janssen WG, Phillips GR (2010) Streamlined embedding of cell monolayers on gridded glass-bottom imaging dishes for correlative light and electron microscopy. Microsc Microanal 16: 747–754

Hirose T, Nakano Y, Nagamatsu Y, Misumi T, Ohta H, Ohshima Y (2003) Cyclic GMP-dependent protein kinase EGL-4 controls body size and lifespan in C elegans. Development 130: 1089–1099

Hosono R (1978) Sterilization and growth inhibition of Caenorhabditis elegans by 5-fluorodeoxyuridine. Exp Gerontol 13: 369–374

Hualin F, Jilong L, Peng D, Weilin J, Daxiang C, 2019. Metabolic wastes are extracellularly disposed by excretosomes, nanotubes and exophers in mouse HT22 cells through an autophagic vesicle clustering mechanism. bioRxiv.

Iida A, Arai HN, Someya Y, Inokuchi M, Onuma TA, Yokoi H, Suzuki T, Hondo E, Sano K (2019) Mother-to-embryo vitellogenin transport in a viviparous teleost. Proc Natl Acad Sci U S A 116: 22359–22365

Kachur T, Ao W, Berger J, Pilgrim D (2004) Maternal UNC-45 is involved in cytokinesis and colocalizes with non-muscle myosin in the early Caenorhabditis elegans embryo. J Cell Sci 117: 5313–5321

Kamath RS, Ahringer J (2003) Genome-wide RNAi screening in Caenorhabditis elegans. Methods 30: 313–321

Melentijevic I, Toth ML, Arnold ML, Guasp RJ, Harinath G, Nguyen KC, Taub D, Parker JA, Neri C, Gabel CV et al (2017) C. elegans neurons jettison protein aggregates and mitochondria under neurotoxic stress. Nature 542: 367–371

Mitchell DH, Stiles JW, Santelli J, Sanadi DR (1979) Synchronous growth and aging of Caenorhabditis elegans in the presence of fluorodeoxyuridine. J Gerontol 34: 28–36

Nelson GA, Lew KK, Ward S (1978) Intersex, a temperature-sensitive mutant of the nematode Caenorhabditis elegans. Dev Biol 66: 386–409

Nicolás-Ávila JA, Lechuga-Vieco AV, Esteban-Martínez L, Sánchez-Díaz M, Díaz-García E, Santiago DJ, Rubio-Ponce A, Li JL, Balachander A, Quintana JA et al (2020) A Network of Macrophages Supports Mitochondrial Homeostasis in the Heart. Cell 183: 94–109.e123

Paul-Gilloteaux P, Heiligenstein X, Belle M, Domart MC, Larijani B, Collinson L, Raposo G, Salamero J (2017) eC-CLEM: flexible multidimensional registration software for correlative microscopies. Nat Methods 14: 102–103

Perez MF, Francesconi M, Hidalgo-Carcedo C, Lehner B (2017) Maternal age generates phenotypic variation in Caenorhabditis elegans. Nature 552: 106–109

Perez MF, Lehner B (2019) Vitellogenins - Yolk Gene Function and Regulation in. Front Physiol 10: 1067

Pokrzywa W, Hoppe T (2013) Chaperoning myosin assembly in muscle formation and aging. Worm 2: e25644

Porta-de-la-Riva M, Fontrodona L, Villanueva A, Cerón J (2012) Basic Caenorhabditis elegans methods: synchronization and observation. J Vis Exp: e4019

Praitis V, Casey E, Collar D, Austin J (2001) Creation of low-copy integrated transgenic lines in Caenorhabditis elegans. Genetics 157: 1217–1226

Redemann S, Schloissnig S, Ernst S, Pozniakowsky A, Ayloo S, Hyman AA, Bringmann H (2011) Codon adaptation-based control of protein expression in C. elegans. Nat Methods 8: 250–252

Van Rompay L, Borghgraef C, Beets I, Caers J, Temmerman L (2015) New genetic regulators question relevance of abundant yolk protein production in C. elegans. Sci Rep 5: 16381

Van Voorhies WA, Fuchs J, Thomas S (2005) The longevity of Caenorhabditis elegans in soil. Biol Lett 1: 247–249

Watanabe S, Punge A, Hollopeter G, Willig KI, Hobson RJ, Davis MW, Hell SW, Jorgensen EM (2011) Protein localization in electron micrographs using fluorescence nanoscopy. Nat Methods 8: 80–84

Zhang Y, Werling U, Edelmann W (2012) SLiCE: a novel bacterial cell extract-based DNA cloning method. Nucleic Acids Res 40: e55

Zhong L, Yuan L, Rao Y, Li Z, Zhang X, Liao T, Xu Y, Dai H (2014) Distribution of vitellogenin in zebrafish (Danio rerio) tissues for biomarker analysis. Aquat Toxicol 149: 1–7

Śliwińska MA, Cały A, Borczyk M, Ziółkowska M, Skonieczna E, Chilimoniuk M, Bernaś T, Giese KP, Radwanska K (2020) Long-term Memory Upscales Volume of Postsynaptic Densities in the Process that Requires Autophosphorylation of αCaMKII. Cereb Cortex 30: 2573–2585

