## Supplemental Table 1 for "Nutritional status and fecundity are synchronised by muscular exopheresis"

| Strain | Genotype | Usage | Reference |
| --- | --- | --- | --- |
| N2 | Wild type | Figure 3E | Brenner 1974 |
| ACH91 | *wacIs6[myo-3p::pas-7::GGGGS Linker-wrmScarlet::unc-54 3’UTR, unc-119(+)], wacIs14[myo-3p::tomm-20_1-50aa::attB5::mGFP::unc-54-3'UTR, unc-119(+)]* | Figure 1A, I; Figure 3F;  Figure 4E | Generated for this study |
| ACH93 | *wacIs1[myo-3p::rpn-5 CAI=0.97::GGGGS Linker-wrmScarlet::unc-54 3’UTR, unc-119(+)], wacIs14[myo-3 promoter::tomm-20_1-50aa::attB5::mGFP::unc-54-3'UTR, unc-119(+)]* | Figure 1A-B, D-H; Figure 2A, D-E; Figure 3A-C, E; Figure 4B-D, G, I; Figure S1A-B;  Figure S2  Figure S3A | Generated for this study |
| ACH199 | *wacIs1[myo-3p::rpn-5 CAI=0.97::GGGGS Linker-wrmScarlet::unc-54 3’UTR, unc-119(+)], vit-2(crg9070[vit-2::gfp]) X* | Figure 1C,  Figure 4F,H; Figure S1C-D  Figure S3B | Generated for this study |
| AGD885 | *rrf-3(b26) II; fem-1(hc17) IV; uthEx633 [myo-3p::GFP]* | Figure 2B-C | Vilchez et al. 2012 |
| CB4088 | *him-5(e1490)V* | Figure 2B | Caenorhabditis Genetics Centre |

**Supplementary Table 1. List of *C. elegans* strains used in this study**
